# Cigarette Smoke Extract (CSE) reduces expression of functional TRPV4 channels in primary human bronchial epithelial cells differentiated at an Air Liquid Interface (ALI) in vitro

**DOI:** 10.64898/2026.05.20.726480

**Authors:** Isabel Müller, Philipp Alt, Thomas Gudermann, Martina Kiefmann, Alexander Dietrich

**Affiliations:** Walther Straub Institute of Pharmacology and Toxicology, Member of the German Center for Lung Research (DZL), Medical Faculty, LMU-Munich, Munich, Germany

**Author notes:** **Corresponding author:** Alexander Dietrich.

**Keywords:** Human bronchial epithelial cells, Ciliated cells, OS-9, Polyubiquitination, Smoke regulated genes, Transient receptor potential vanilloid 4

## Abstract

Primary human bronchial epithelial cells (pHBECs) of the airways of smokers are chronically exposed to cigarette smoke, which may induce chronic obstructive pulmonary disease (COPD) ranked fourth among the most common global causes of death. Using an established protocol for differentiation of pHBECs to a pseudostratified epithelium at an air liquid interface (ALI), we analyzed functional expression of transient receptor potential vanilloid 4 (TRPV4) proteins after application of cigarette smoke extract (CSE), which upregulated seven smoke exposure regulated genes (SERGs). TRPV4 protein expression in the plasma membrane and localization next to the cilia of ciliated cells was reduced, while cell barrier function was not altered after chronic exposure to CSE for 28 days compared to untreated control cells. Accordingly, TRPV4-mediated Ca^2+^ influx was blocked in pHBECs after CSE exposure. Moreover, Os-9 protein, which after binding mediates protection from degradation of TRPV4 protein by polyubiquitination, was significantly less expressed in pHBECs upon CSE exposure. Most interestingly, overexpression of OS-9 in pHBECs rescued reduced TRPV4 protein levels induced by CSE. Our study identifies a novel molecular mechanism of toxicity by CSE interfering with TRPV4 and OS-9 expression in pHBECs, which may blaze the trail for new therapeutic options in COPD.

## Introduction

Smoking induces 8 out of 10 common causes of deaths worldwide every year (Jha et al. 2015) mostly by inducing chronic obstructive pulmonary disease (COPD) (Calverley and Walker 2023). Primary human bronchial epithelial cells (pHBECs) are one of the first targets of cigarette smoke (CS), which consists of approximately 7000 compounds in the gas and particle phase. pHBECs serve different specific functions to reduce toxicity of CS. Basal cells are able to differentiate to all cell types of the epithelium, while club cells secret surfactant, are progenitors for ciliated and goblet cells as well as dedifferentiate to basal cells after injury (Wu et al. 2022). Moreover, ciliated and goblet cells are the key cells for mucociliary clearance of particles in the tracheobronchial region of the airways. While goblet cells secret mucus to trap them, ciliary movement by ciliated cells carry them upwards and a cough reflex removes them from the airways (reviewed in (Hiemstra et al. 2015)). However, excess mucus production, chronic cough and inflammation of the lower airways are also typical symptoms of COPD (Calverley and Walker 2023) currently ranked fourth of the most common global causes of death.

The fourth member of the transient receptor potential vanilloid (TRPV4) family is prominently expressed in the lung endothelium (Simmons et al. 2018) as well as in the alveolar (Weber et al. 2020) and tracheobronchial epithelium (Alt et al. 2026). Like most TRP channels, TRPV4 harbors an invariant sequence, the TRP box (containing the amino acid sequence: EWKFAR), in its intracellular C-terminal tail as well as ankyrin repeats in the intracellular N-terminus. The protein is composed of six membrane-spanning helices (S1-6), and a presumed pore-forming loop between S5 and S6 (Nilius and Szallasi 2014). Four of these TRPV4 monomers assemble in homo-tetrameric complex with a functional common pore domain, which is permeable for cations like Ca^2+^ and Na^+^ (Hellwig et al. 2005). TRPV4 proteins were originally identified as a sensor of extracellular osmolarity (Liedtke et al. 2000; Strotmann et al. 2000), but may also serve as mechano-sensors because they are activated by membrane and shear stretch as well as by viscous loading (Goldenberg et al. 2015a). Most interestingly, TRPV4 channels regulate ciliary beat function for mucociliary clearance by ciliated cells (Lorenzo et al. 2008) and are involved in the progression of pulmonary hypertension (Goldenberg et al. 2015b; Xia et al. 2014) and lung fibrosis (Rahaman et al. 2014). We recently demonstrated that TRPV4 channels mediate surfactant protein A and D secretion in club cells and contribute to basal cell differentiation in ciliated cells using a human air liquid interface (ALI) model in vitro (Alt et al. 2026). Here, we use this ALI model to analyze TRPV4 expression and function in airway epithelia cells after chronic application of cigarette smoke extract (CSE). CSE reduced TRPV4 expression in the lung epithelium and localization of the channel in near proximity to cilia in ciliated cells. Moreover, OS-9 upregulation rescued TRPV4 protein from polyubiquitination and subsequent degradation. Therefore, TRPV4 channels and its interaction partner OS-9 may serve as future therapeutic target to reduce toxic effects of CSE in the airway epithelium.

## Materials and Methods

### Generation of Cigarette Smoke Extract (CSE)

The production of CSE was performed as described previously (Mastalerz et al. 2022; Schamberger et al. 2015). Six research-grade cigarettes with filters (3R4F, Kentucky Tobacco Research and Development Center at the University of Kentucky; Lexington, KY) were bubbled in a device (Figure S1A) through 100 mL PneumaCult™-ALI medium (Stemcell Technologies, Canada, #05001) with a burning time of 4 minutes each. The received CSE is designated as 100 % CSE, sterile filtered and the pH value was adjusted if necessary. Aliquots were stored at -80 °C and diluted to a final concentration of 5 % CSE freshly before use. CSE application started with the airlift (d0) at the basolateral compartment and continued throughout the whole differentiation period (28 days).

### Human ALI model

Primary human bronchial epithelial cells (pHBECs) from healthy donors were obtained from Lonza (Basel, Switzerland, CC#2540S, see Supplemental Table S1 for donor information) and processed as described (Alt et al. 2026). In brief, cells were expanded for two passages in PneumaCult™Ex Plus medium (Stemcell Technologies, Canada, #05040) or PneumoCult NGEX medium (Stemcell Technologies, Canada. #100-1505). Afterwards cells were seeded onto collagen IV-coated (Sigma Aldrich, Germany, #C6745) 12-well Transwell-inserts (Corning, USA, #3460) with a density of 120000 cells/insert. Cells were cultured at 37 °C and 5 % CO2 and the air-lift was performed 2-3 days after seeding when confluency was reached. The medium at the basal compartment was changed to PneumaCult™ ALI-medium (Stemcell Technologies, Canada. #05001) or 5% CSE diluted in PneumaCult™ ALI-medium and the medium at the apical compartment was removed. From then on medium change was performed every two days. Cells were washed with prewarmed DPBS once a week to remove secreted mucus beginning with the second week after the air lift and wash-offs were collected for further analysis. Depending on the experimental setup, cells were cultured for 7, 14, 21, and/or 28 days.

### Cell viability Assay

To assess the cytotoxicity of the generated CSE, a lactate dehydrogenase (LDH) colorimetric assay (F. Hoffmann-La Roche AG, Switzerland, #11644793001, version 10) was performed as described before (Schamberger et al. 2015). LDH values were quantified from the basolateral medium after application of different concentration of CSE. As positive control, cells were lysed in 2% Triton-X 100 diluted in cell specific medium. The quantification was performed according to the manufacturer’s protocol

### Immunofluorescence staining and confocal imaging

Differentiated pHBECs were washed with cold PBS, fixed in 4% PFA/PBS (15 min at RT) on inserts and then washed thrice with PBS. Cells were permeabilized for 15-60 min (depending on time point of differentiation) with 0.5 % Triton X100 solution in PBS and then washed 3 × 5 min with DPBS. Blocking solution containing 4 % BSA in DPBS was applied for 1 h at RT and inserts were incubated at 4 °C overnight with primary antibodies (diluted in 2 % BSA in DPBS; club cell marker CC10, Santa Cruz Biotech, Texas, USA, #sc-365992AF488, 1:200; goblet cell marker MUC5AC, Abcam, Cambridge, UK, #ab218714, 1:200; ciliated cell marker acetyl-Tubulin, Abcam, Cambridge, UK, #ab218591). The next day, inserts were washed 3 × 5 min with 0.1 % BSA in DPBS. Nuclei staining was performed using DAPI solution (0.1 mg/L in DPBS) for 15 min at RT. Inserts were washed again 2 × 5 min with 0.1 % BSA in DPBS and finally 2 × 5 min with DPBS. Inserts were cut out, mounted on glass cover slides (Epredia, New Erie Scientific, Netherlands, #J1800AMNZ) with mounting medium (ProLong™ Glass Antifade Mountant, Thermo Fisher Scientific, USA, #P36980). Slides were stored at 4 °C for imaging. We used fluorescent dye coupled antibodies for the experiments with different wavelengths (488 nm for p63 and CC10, 555 nm for MUC5AC, 647 nm for acetyl-tubulin). Confocal images were taken as z-stacks with a Zeiss LSM 880 microscope using ZEN Black software (version 2.3). Image processing was performed by using ZEN Blue software (version 3.4, Carl Zeiss, Jena, Germany) and Fiji software (Image J v. 1.53c, Wayne Rasband, NIH, USA). Stacks were separated with respective channels (405 nm, 488 nm, 555 nm, 647 nm) and composed 2D-images were generated. For analysis, every insert was imaged 5 times on different areas across the insert with a 20 x air objective, 40 x oil objective, 63 x water objective or 100 x oil objective. For each time point, at least two technical replicates for each condition and experiment were performed.

### Permeability Assay

To verify the epithelial integrity, a permeability assay was performed at different time points of the differentiation. Fluoresceinisothiocyanat (FITC)–Dextran (Sigma Aldrich, USA, #FD20S) was solved in HBSS without phenol red (containing Mg^2+^ and Ca^2+^) to a final concentration of 1 mg/mL. Afterwards, cells were washed with cold DPBS on both sites, 250µL FITC–Dextran solution were added to the apical compartment, while 800 µL prewarmed HBSS were added to the basolateral compartment. Cells were kept in the dark for 20 minutes at room temperature. Afterwards, 100 µL of the apical and basolateral solution were transferred into a black 96-well microplate with clear bottom (Corning® Inc., USA, #CLS3603) to measure the fluorescence signal at 490/520 nm.

### Ca^2+^ Imaging of fully differentiated pHBECs

Primary HBECs were differentiated on 12mm collagen-coated Transwell™ inserts for 28 days. On the day of measurement, the fully differentiated epithelium (day 28) was loaded with 10μM Fura-2 AM (Invitrogen™, Thermo Fisher Scientific, #F1221) in HBSS (containing Ca^2+^ and Mg^2+^) for 40 min at 37 °C. After incubation, inserts were washed with HBSS, cut out and placed into a quick-change chamber (Warner Instruments, Holliston, USA) with 150 μL of HBSS. Imaging was performed with a 20 x air objective of a Leica DM98 fluorescence microscope. Intracellular Ca^2+^-concentration ([Ca^2+^]_i_) was measured after the application of TRPV4-activator (GSK10160970A, Merck, # 5.30533, 100 μM or 200 μM or 500 µM) with or without using TRPV4-blocker (HC067047, Merck, # 616521). Cells were excited at 340 and 380 nm and fluorescence emissions were recorded at 510 nm. Emission ratios at wave lengths of 340 nm and 380 nm were calculated.

### SDS-PAGE and Western Blot analysis

Expression of proteins was determined by Western blot analysis, as previously described (Hofmann et al. 2017). Cells from two inserts were lysed each with 150 μL RIPA buffer (containing phosphatase and protease inhibitors) and placed on ice for 30 min. Protein concentration was measured with the Pierce BCA Protein Assay Kit (Thermo Fisher Scientific, #23225) according to manufacturer’s protocol. The protein samples were mixed with 2 x (Sigma Aldrich, USA, #S3401-10VL) or 5 x laemmli buffer (3 mL 2.6 M TRIS/HCl pH 6.8, 10 mL glycerol, 2 g SDS, 2 mg bromophenol blue, 5 mL β-mercaptoethanol) and heated for 10 min at 95 °C. Then, samples with 10, 15, 20 or 25 μg of total protein were loaded onto an SDS-PAGE gel (stacking gel 4% polyacrylamide, separating gel 10 % polyacrylamide) and electrophoresis was run for 30 min at 70 V at separating gel and then for 100 min at 100 V. Proteins then were transferred from the gel to a Roti®-PVDF membrane (Carl Roth, Karlsruhe, Germany, #T830.1) in a wet-transfer system from BioRad (Feldkirchen, Germany) with 90 mA for 16 h at 4 °C. Blocking was performed using 5 % low-fat milk powder (Carl Roth, #T145.2) in TBS-T (0.1 % TWEEN 20 in TBS buffer) for 1 h at RT. All antibodies were diluted according to supplier’s datasheet in blocking solution (TRPV4-antibody, Merck, Germany, #MABS466, 1:10.000; OS-9-antibody, Abcam, UK, #ab109510, 1:1.000; anti acetyl-tubulin, Abcam, UK, #ab179484, 1:2.000; anti-ß-actin-POX, Merck, Germany, #A3854, 1:10.000; anti-mouse-IgG-HRP, Cell Signaling, Danvers, Massachusetts, USA, #7076, 1:2.000; anti-rabbit-IgG-POX, Sigma Aldrich, USA, #A6151, 1:10.000). Membranes were incubated in primary antibodies solution overnight (16 h) at 4 °C, washed 3 × 5 min with TBS-T and incubated again with peroxidase-conjugated secondary antibody solutions for 2 h at RT. After washing again 3 × 5 min with TBS-T, membranes were incubated with SuperSignal West Pico, Femto or Atto (Thermo Fisher Scientific, #34577) sensitivity substrates and imaged using an Odyssey-Fc unit (Licor, Lincoln, NE, USA).

### Quantitative reverse transcription PCR (qRT-PCR)

Gene-expression of collected samples was measured with quantitative Reverse-Transcriptase Polymerase Chain Reaction (qRT-PCR). Therefore, cells were lysed and RNA was isolated via RNeasy® Plus Mini-Isolation Kit (Qiagen, Germany, #74136) according to the manufacturer’s protocol. Then, 500 ng of RNA was transcribed into cDNA with RevertAid H Minus First Strand cDNA Synthesis Kit (Thermo Fisher Scientific, USA, #K1632). Quantitative qPCR was performed using dilution of cDNA with specific oligonucleotides for genes of interest (sequence see Table S2) and ABsolute qPCR SYBR Green Mix (Thermo Fisher Scientific, USA, #AB-1285/B) in a 96 well-plate and was analyzed with a LightCycler480 from Roche. Received Ct-values were taken for calculation of expression of the respective genes normalized to ribosomal protein lateral stalk subunit P0 (RPLP0) as housekeeping gene. Normalized values of samples were then referred to ctrl samples for detection of possible differences between ctrl and CSE-treated samples.

### Heterologous overexpression of OS-9 in differentiated pHBECs

To verify OS-9’s role in the polyubiquitination of TRPV4, pHBECs differentiated under CSE exposure were transfected with a plasmid containing OS-9-eGFP in the fourth week of the differentiation. The transfection with Lipofectamine 3000 Reagent (Invitrogen™, Thermo Fisher Scientific, California, USA, #3000150) was performed according to the manufacturer’s protocol using 3 µg of DNA per well. Transfection mix was added to the apical site and incubated for 2 days. Cells were lysed in RIPA buffer for further analysis.

### Proximity Ligation Assay (PLA)

To investigate the interaction between OS-9 and TRPV4, fully differentiated pHBECs were washed with cold DPBS and fixed in 4% PFA solution as described above. The cells were treated according to the steps of the manufacturer’s protocol. In brief, cells were blocked in blocking solution and incubated with primary antibodies (from different hosts) against OS-9 and TRPV4. Then, PLA-probes and ligation buffer for rolling circle amplification were added. After amplification step and incubation with fluorescent-labeled oligonucleotide probes (complementary to the amplified product), cells were washed and mounted in DAPI-containing mounting-medium (Merck, Germany, anti-mouse MINUS #DUO92004-30RXN, anti-rabbit PLUS #DUO92002-30RXN). After solidification, the cells were imaged at 595 nm with a confocal microscope (LSM 880, Zeiss, Germany). The interaction signal was quantified with Fiji software (Image J v. 1.53c, Wayne Rasband, NIH, USA) by measuring the integrated density of single cells.

### Ubiquitin Enrichment Analysis

The ubiquitination status of TRPV4 was analyzed with a Pierce ™ Ubiquitin Enrichment Kit (Thermo Scientific™, USA, #89899). Fully differentiated pHBECs were lysed as described above with 150 µL RIPA buffer. The obtained cell lysate was centrifuged at 15.000 x g for 15 minutes at 4 °C and the protein containing supernatant was transferred into a clean and prechilled tube for further analysis. The protein content was determined with the Pierce™ BCA Protein Assay Kit. 150 µg of total protein was applied to a column with polyubiquitination affinity resin and eluted according to the manufacturer’s protocol. Analysis of the purified samples was performed via SDS PAGE and Western Blot as described above. Samples were volume-normalized to the unprocessed fraction and 15 µg of the unpurified sample served as loading control.

### Statistical analysis

For statistical analysis, GraphPad Prism 10 software (GraphPad Software, San Diego, USA) was used. Significant differences are indicated by asterisks. All data are represented as means + SEM.

## Results

### Experimental setup of a human air liquid interface (ALI) for detection of acute and chronic airway toxicity by cigarette smoke extract (CSE)

The experimental setup and the timeline for the detection of acute airway toxicity is outlined in Figure 1A. Primary human bronchial epithelial cells (pHBECs) from healthy human donors (non-smokers, no alcohol abuse, see Table S1) were expanded and seeded on inserts. After the airlift, differentiation of a pseudostratified epithelium occurred in 28 days as already described (Mastalerz et al. 2022; Schamberger et al. 2015). Different concentrations of the produced cigarette smoke extract (Figure S1B) were applied to the lower compartment chronically after air lift and acute as well as chronic toxicity were quantified by applying a lactate dehydrogenase (LDH) assay. Cell viability was not significant different after application of 5, 15 and 50 % of CSE compared to the 0% control values. 100% CSE however induced an acute significant drop in cell viability (Figure S1B). We and others choose 5% CSE to mimic chronic exposure to CSE, which was applied to the basal compartment after airlift (Mastalerz et al. 2022). As described, this technique was identified as the best method and resulted in a successful up-regulation of mRNA expression from seven smoke regulated genes (SERGs) like in chronic smokers (Figure S2 (Mastalerz et al. 2022)).

**Fig. 1.**
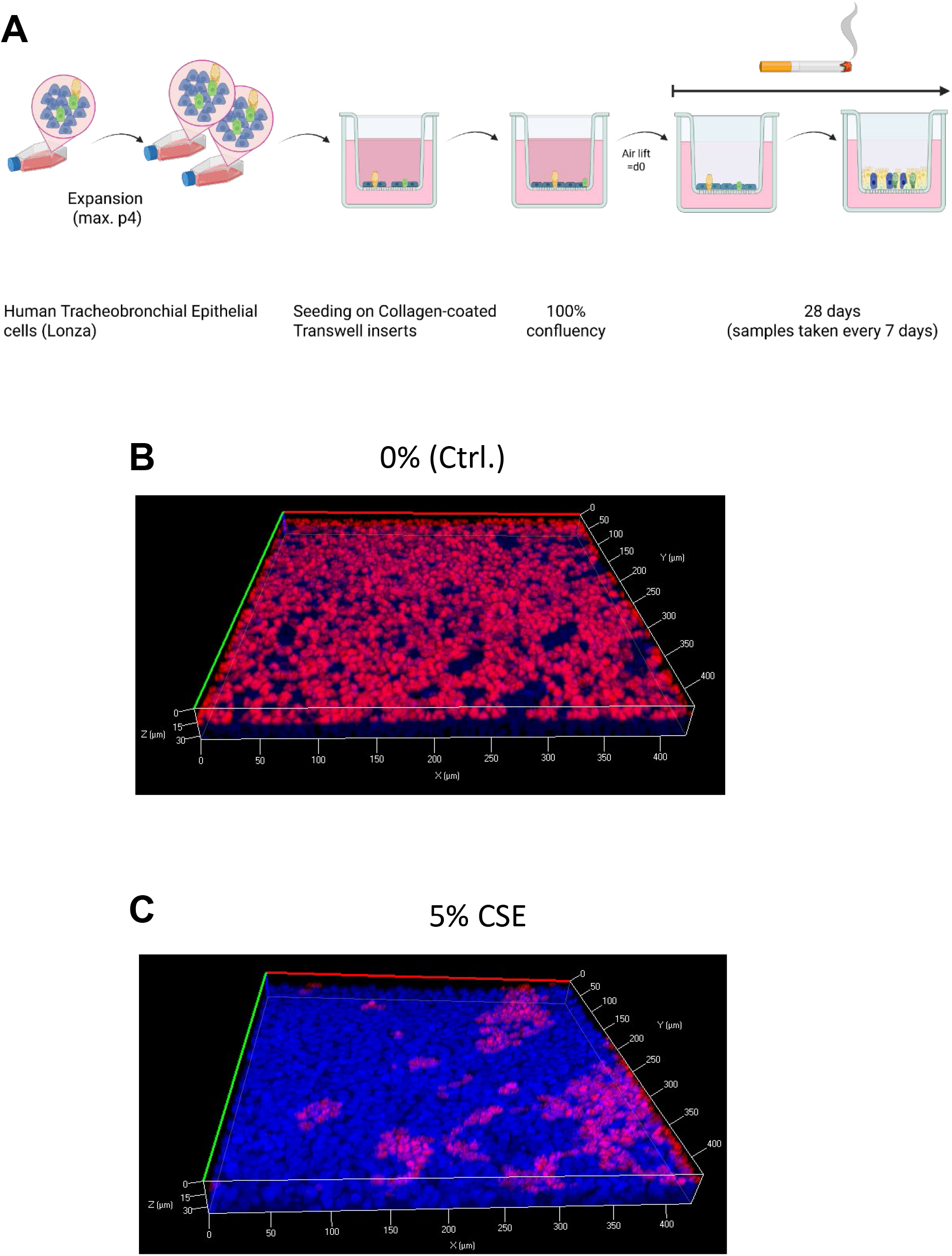
Application of cigarette smoke extract (CSE) to primary human bronchial epithelial cells (pHBECs) differentiated at the air liquid interface (ALI) results in loss of ciliated cells in the pseudostratified epithelium. **A** Experimental set up of the ALI differentiation model. After expansion of pHBECs, cells were seeded to filter inserts and after confluency and air lift differentiated to a pseudostratified epithelium. CSE (5%) was added to the medium in the lower compartement after airlift for a chronic exposure of 28 days. Medium was exchange every second day. No CSE was added to control cells. **B** Representative confocal image of the differentiated pseudostratified epithelium without CSE exposure (0%, Ctrl.). Ciliated cells were identified by staining with fluorescent coupled antibodies directed against acetylated tubulin proteins (red). All cell nuclei were stained with DAPI (blue). **C** Representative confocal image of the differentiated pseudostratified epithelium after chronict CSE exposure (5% CSE). Ciliated cells were identified by staining with fluorescent coupled antibodies directed against acetylated tubulin proteins (red). All cell nuclei were stained with DAPI (blue).

### Chronic application of CSE reduced number of ciliated cells and TRPV4 channels in an ALI model of the bronchial epithelium

Chronic application of CSE resulted in a significant reduction of ciliated cells compared to control cells as already described (Figure 1B, C,(Schamberger et al. 2015)), which was also apparent in a reduced expression of ciliated tubulin (Figure S3A, B). Although ciliated cells as well as goblet and club cells were reduced after CSE exposure (Figure S3C), loss of differentiated cells did not result in a higher permeability of the airway epithelium as detected in a fluorescence permeability assay (Figure S4). Along this line, the numbers of total cells detected by staining of nuclei with 4’, 6’ diamidino-2-phenylindole dihydrochloride (DAPI) was obviously not different (Figure 1B, C and Figure S3C). As we recently identified a lower number of ciliated cells after down-regulation of TRPV4 protein in a human ALI model (Alt et al. 2026), we next analyzed TRPV4 expression after CSE exposure. Quantitative Western Blotting of cell lysates of the bronchial epithelium at different time points of differentiation revealed reduced numbers of TRPV4 protein after application of CSE compared to control cells (Figure 2A, B). Apparent was also a significantly different localization of TRPV4 protein in airway epithelial cells (Figure 2C, D). While in control cells the majority of TRV4 protein was located close to the acetylated tubulin of cilia in ciliated cells and was evenly distributed in lower cell layers (Figure 2C), application of CSE resulted mainly in expression of TRPV4 protein in a cell layer directly under the cilia (Figure 2D). To reproduce these results on a functional level, we quantified Ca^2+^ influx through TRPV4 channels in epithelial cells after application of a TRPV4 activator. Indeed, an increased Ca^2+^ influx after application of a TRPV4 agonist was only detected in control cells, but not in cells treated with CSE (Figure 3A-C). Surprisingly, mRNA expression of TRPV4 proteins was not reduced, but significantly increased in epithelial cells after application CSE (Figure S5), pointing to a post-translational mechanism of protein loss by CSE.

**Fig. 2.**
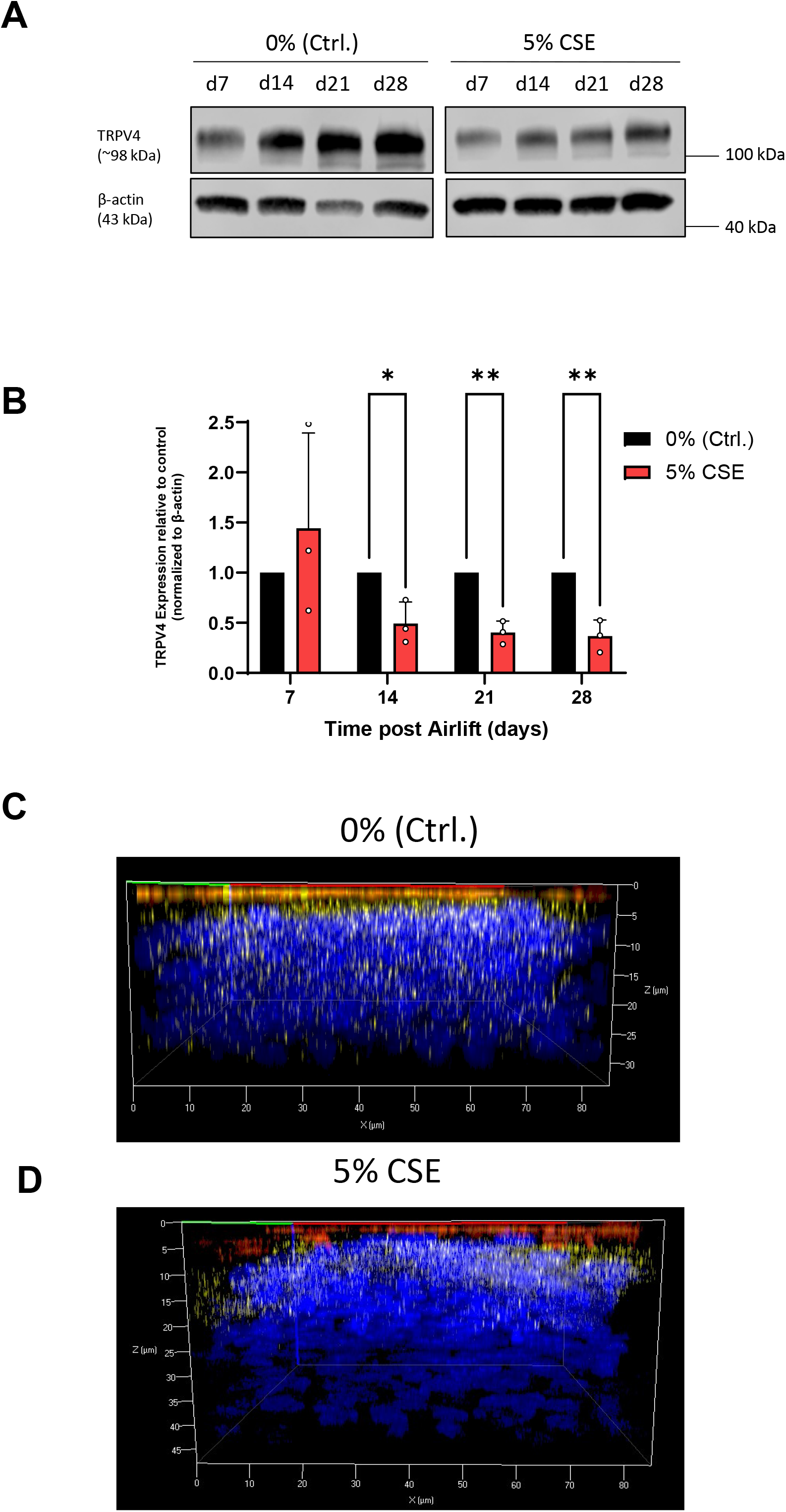
CSE induces loss and changed localisation of TRPV4 protein in pHBECS. **A** Representative Western Blot processed with a specfic TRPV4 antibody from cell lysates of pHBECs differentiated for the indicated time points at an ALI with (5% CSE) or without application of CSE (0% (Ctrl)). Beta actin was used as loading control. **B** Quantification of TRPV4 protein expression by Western Blotting of pHBECs lysates incubated with (5% CSE, red bars) or without CSE (0% (Ctrl.), black bars) and differentiated for the indicated time points at an ALI. Values are shown relative to controls and normalized to β-actin expression. All data represent means + SEM from at least 3 experiments. Significance between means was analyzed using multiple unpaired Student’s t-test and is indicated as ** for p<0.01 and * for p<0.05. **C** Confocal image showing a cross section through an ALI culture differentiated for 28 days under control conditions (0% (Ctrl.)). **D** Confocal image showing a cross section through an ALI culture differentiated for 28 days after application of CSE (5% CSE).

**Fig. 3.**
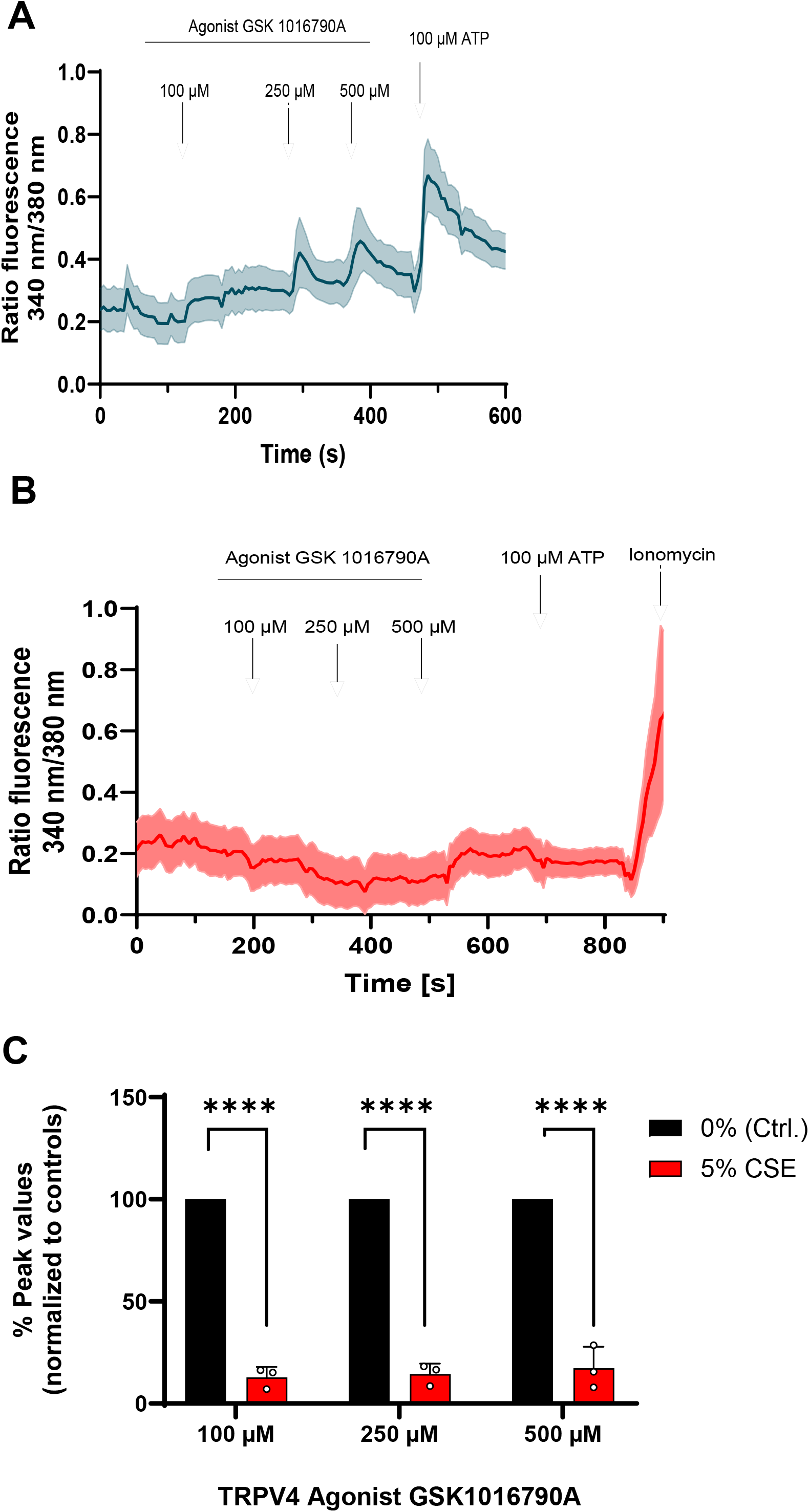
Ca^2+^ imaging reveal a loss of increases in intracellular Ca^2+^ concentrations ([Ca^2+^]_i_) in differentiated pHBECs at an ALI after chronic preincubation with CSE and application of a TRPV4 activator. **A** [Ca^2+^]_i_ as fluorecence ratios in differentiated pHBECs without preincubation of CSE. A TRPV4 agonist (GSK 1016790A) was added at the indicated time points in specified concentrations with a final application of ATP (100 mM). **B** [Ca^2+^]_i_ as fluorescence ratios in differentiated pHBECs with preincubation of CSE (5%). A TRPV4 agonist (GSK 1016790A) was added at the indicated time points in specified concentrations with a final application of ATP (100 mM) and Ionomycin. Data present means +/-SEM for at least 25 cells in one representative analysis out of three experiments. **C** Quantification of peak values from increases of [Ca^2+^]_i_ in differentiated pHBECs with (5% CSE, red bars) or without (0% (Ctrl.), black bars) induced by a TRPV4 activator. Data present means + SEM for at least three Ca^2+^ imaging experiments. Significance between means was analyzed using multiple unpaired Student’s t-test and is indicated as **** for p<0.0001.

### OS-9 expression and its interaction with TRPV4 channels is reduced after CSE exposure

It was reported that a protein, OS-9, binds directly to nascent TRPV4 monomers and blocks its polyubiquitination and degradation (Wang et al. 2007). Therefore, we quantified OS-9 expression in lysates of bronchial epithelial cells and identified a significant loss of OS-9 expression after application of CSE in comparison to control cells (Figure 4A, B). Therefore, we also quantified OS-9 – TRPV4 interaction using a proximity ligation assay. OS-9 – TRPV4 interaction was also significantly reduced after application of 5% CSE compared to cells not exposed to CSE (Figure 4C, D).

**Fig. 4.**
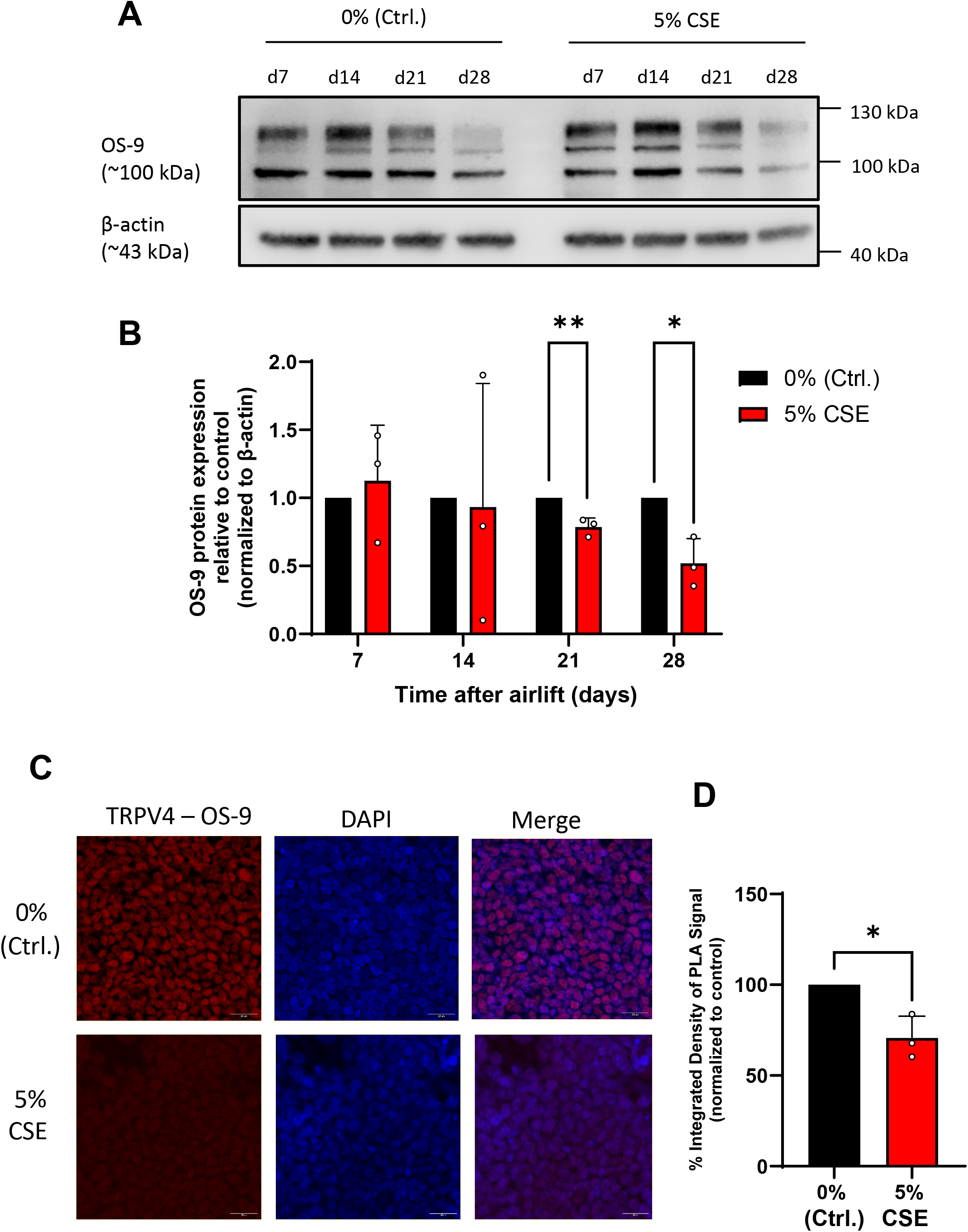
OS-9 expression and interaction with TRPV4 channels is reduced in pHBECs differentiated at an ALI by chronic application of CSE. **A** Representative Western Blot processed with a specfic OS-9 antibody from cell lysates of pHBECs differentiated for the indicated time points at an ALI with (5% CSE) or without application of CSE (0% (Ctrl)). Beta actin was used as loading control. **B** Quantification of OS-9 protein expression by Western Blotting of pHBECs lysates incubated with (5% CSE, red bars) or without CSE (0% (Ctrl., black bars) and differentiated for the indicated time points at an ALI. All values are shown relative to controls and were normalized to β-actin expression. Data represent means + SEM from at least 3 experiments. Significance between means was analyzed using multiple unpaired Student’s t-test and is indicated as ** for p<0.01 and * for p<0.05. **C** Confocal images of pHBECs differentiated for 28 days with (upper panels) or without preincubation of CSE (lower panels) after application of a Proximity Ligation Assay (PLA). Direct TRPV4-OS-9 interactions are indicated by red signals (left panels), nuclei were stained with DAPI (middle panels) and merged images (right panels) are depicted. **D** Quantification of TRPV4-OS-9 interactions in differentiated pHEBCs with (5% CSE, red bar) or without preincubation with CSE (0%, (Ctrl.), black bar) identified by a PLA. Data present means + SEM of 3 experiments. Significance between means was analyzed using unpaired Student’s t-test and is indicated as * for p<0.05.

### CSE-induced polyubiquitination of TRPV4 is reduced after over-expression of OS-9 in airway epithelial cells

To identify functional interaction of OS-9 and TRPV4, we quantified polyubiquitination of TRPV4 protein in bronchial epithelial cells with or without CSE exposure. While CSE application gave rise to more polyubiquitinatied TRPV4 protein eluted from the affinity column, very little ubiquitinated TRPV4 protein was eluted in control cells (Figure 5A). Moreover, we were able to overexpress OS-9 protein in pHBECs by transfection with a plasmid coding for the protein in a rescue experiment (Figure 5B). Most interestingly, overexpression of OS-9 resulted in more TRPV4 expression, but less ubiquitinated TRPV4 protein in pHBECs after CSE application (Figure 5B, C). Therefore, reduced OS-9 protein levels and subsequent polyubiquitination of TRPV4 after CSE exposure are at least in part responsible for a loss of functional TRPV4 channels in human bronchial epithelial cells differentiated at an air liquid interface.

**Fig. 5.**
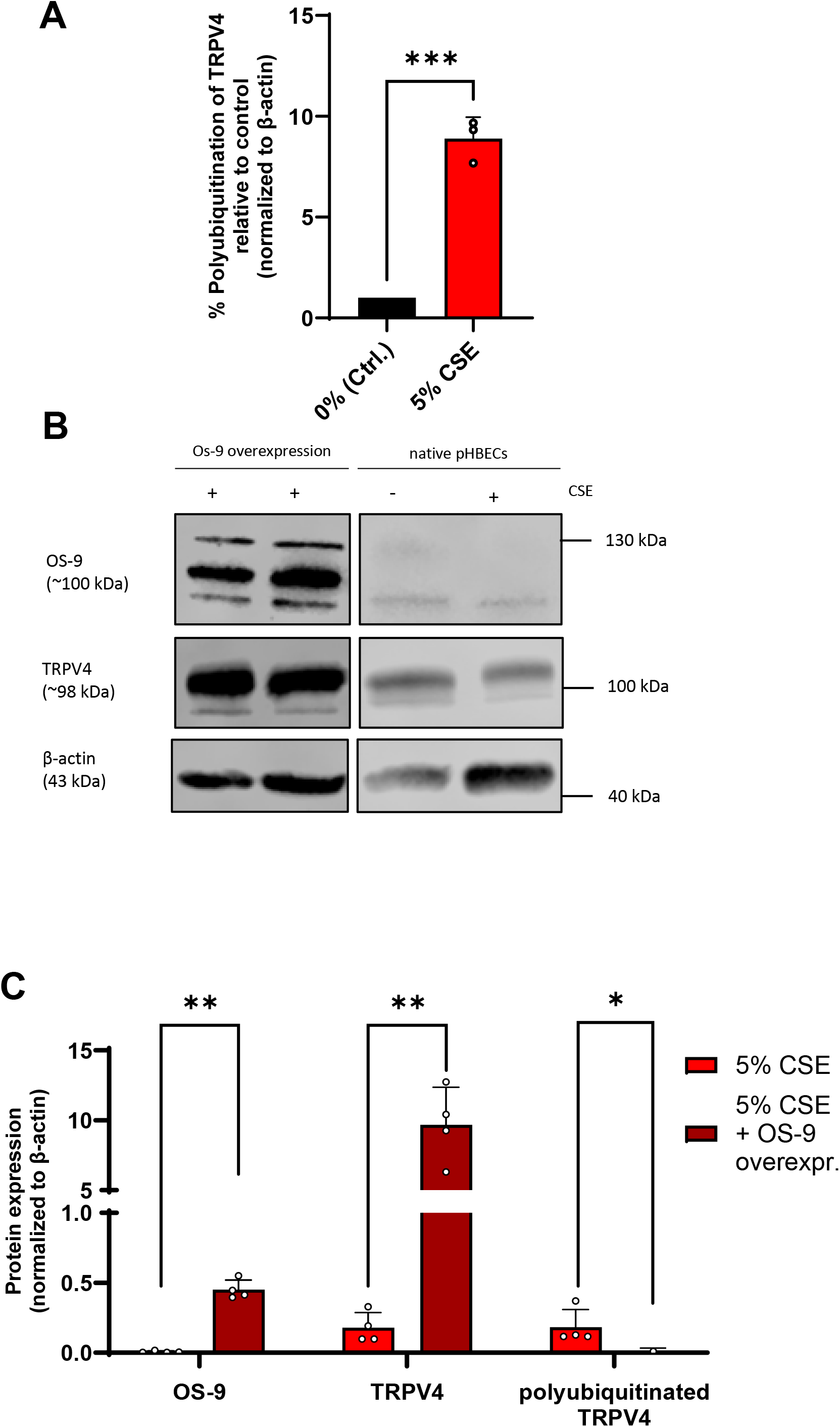
TRPV4 protein expression is reduced and polyubiquitination is increased after application of CSE, while over-expression of OS-9 rescues TRPV4 in pHBECs. **A** Polyubiquitination of TRPV4 proteins in differentiated pHBECs with (5% CSE, red bar) or without preincubation of CSE (0 (Ctrl.), black bar). Data are shown relative to the control and normalized to β-actin. **B** Representative Western Blots showing over expression of OS-9 protein in differentiated pHBECs after transfection with an OS-9 expression plasmid processed with a specfic OS-9 antibody (upper left panels) in comparison to native differentiated pHBECs (upper right panels) with (+) or without (-) chronic preincubation with CSE. TRPV4 protein expression was detected by a specfic TRPV4 antibody after OS-9 over-expression (middle left panels) in comparison to native pHBECs (middle right panels) with (+) or without (-) preincubation with CSE. Beta actin was used as loading control (lower panels). **C** Quantification of OS-9, TRPV4 and polyubiqititinated TRPV4 protein in differentiated pHEBECs incubated with CSE and with (5% CSE, dark red bars) or without over-expression of OS-9 protein (red bars) in pHBECs differentiated for 28 days at ALI. Values are shown relative to controls and were normalized to β-actin expression. Data represent means + SEM from at least 4 experiments. Significance between means was analyzed using multiple unpaired Student’s t-test and is indicated as ** for p<0.01 and * for p<0.05.

## Discussion

Differentiation of pHBECs at the ALI is a powerfull method to study cell polarization and permeabilty, as well as bronchial inflammation (reviewed in (Silva et al. 2023). We and others used the system to study toxicity and their underlying molecular mechanisms (Chakraborty et al. 2025; Schamberger et al. 2015) and applied CSE to the basal compartment of the human ALI model (Figure 1A), as only this techniques warranted an upregulation of all seven smoke regulated genes (SERGs) like in chronic smokers (Figure S2 (Mastalerz et al. 2022)). Chronic application of 5% CSE produced with a standard method from research cigarettes (Figure S1A), resulted in a loss of ciliated cells and their protein marker acetylated tubulin in the finally differentiated pseudostratified epithelium (Figure 1B, C, Figure S3A, B) as described (Schamberger et al. 2015), without loosing cell barrier function (Figure S4). As whole cell numbers identified by staining of nuclei with DAPI did not change signficantly (Figure 1B, C, Figure S3C), a block in ciliated cell differentiation with a survival of undifferentiated cells in the pseudostratified epithelium is the most likely reason for the maintance of barrier function. Loss of ciliated cells in chronic smokers will reduce mucociliary clearance for removal of pathogens and foreign materials, which will result in excess mucus production and chronic cough, two hallmarks of COPD (reviewed in (Hattab et al. 2016)). In the light of our recent publication demonstrating that TRPV4 down-regulation in a human ALI model also resulted in loss of ciliated cells (Alt et al. 2026), we identified reduced TRPV4 protein levels after chronic exposure to CSE (Figure 2A, B). We also identified a clear colocalization of TRPV4 channels with cilia of the airway epithelium, which was almost completely lost after application of CSE (Figure 2C, D). This colocalization was also reported in wild-type but not in TRPV4-deficient (TRPV4-/-) mice (Lorenzo et al. 2008). But in clear contrast to TRPV4-/-mice, where only ciliar beating frequencies were reduced (Lorenzo et al. 2008), we identified a significant loss of ciliated cells and TRPV4 protein during pHBEC differentiation under chronic CSE exposure in our translational approach. To provide additional evidence for the reduction of functional TRPV4 channels in pHBECs, we quantified Ca^2+^ influx induced by a TRPV4 specfic activator (GSK 1016790A, (Thorneloe et al. 2008; Willette et al. 2008)) in pHBECs, which was almost absent after CSE exposure (Figure 3A-C). As OS-9 was reported to bind to TRPV4 proteins and protect them from degradation (Wang et al. 2007), we also quantified OS-9 protein levels and detected lower levels and less interaction with TRPV4 protein by CSE application (Figure 4A-D). Polyubiquitination of TRPV4, which results in degradation was also significantly increased after CSE exposure (Figure 5A, B). A succesfull up-regulation of OS-9 protein after transfection was however able to rescue TRPV4 protein levels in pHBECs after CSE exposure (Figure 5C).

In summary, our results demonstrate a clear correlation of reduced TRPV4 expression with loss of ciliated cells after CSE exposure, which was confirmed by our recently identifed decrease of ciliated cells after down-regulation of TRPV4 protein in pHBECs differentiated at the ALI (Alt et al. 2026). As mucociliary clearance is reduced in heavy smokers and induces excess mucus production and chronic cough, TRPV4 and its interaction partner OS-9 might serve as therapeutical targets to reduce these typical symptoms of COPD in the future.

## Abbreviations

ALI: Air Liquid Interface
[Ca^2+^]_i_: intracellular Ca^2+^ concentration
CSE: Cigarette Smoke Extract
pHBECs: primary Human bronchial epithelial cells
SERGs: Smoke exposure regulated genes
TRPV4: Transient receptor potential vanilloid 4

## Acknowledgements

The authors thank Benedikt Kirmayer and Bastian Tschache for excellent technical assistance.

## Author contributions

IM and AD designed the study. IM performed research and collected data. IM, PA, TG, MK and AD analyzed data. AD wrote the first draft and all authors commented on the manuscript. All authors read and approved the final manuscript.

## Ethical standards

Ethical statements are available from the supplier of the pHBECs (Lonza, Basel, Switzerland).

## Supporting information

**Table S1.**
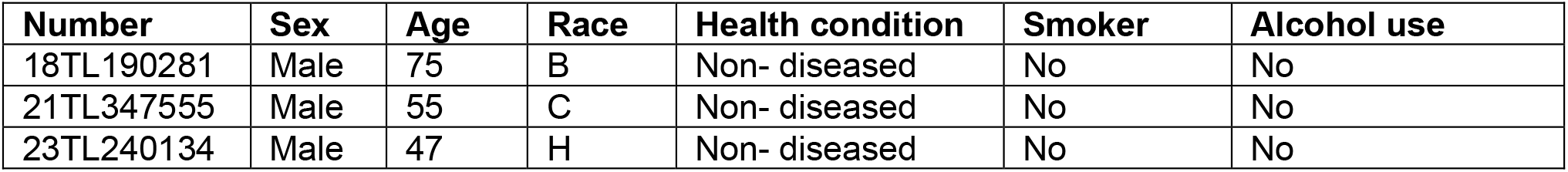
List of human donors of primary human bronchial epithelial cells (pHBECs) differentiated at an ALI:

| Number | Sex | Age | Race | Health condition | Smoker | Alcohol use |
| --- | --- | --- | --- | --- | --- | --- |
| 18TL190281 | Male | 75 | B | Non- diseased | No | No |
| 21TL347555 | Male | 55 | C | Non- diseased | No | No |
| 23TL240134 | Male | 47 | H | Non- diseased | No | No |

**Table S2:**
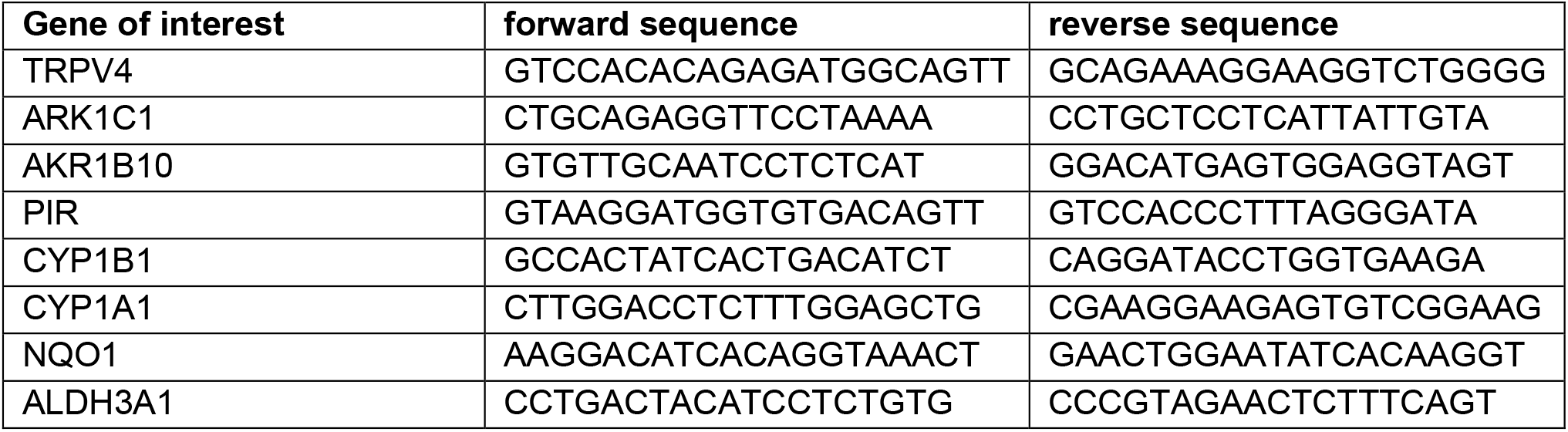
List of oligonucleotide sequences, used for qRT-PCR.

**Fig. S1.**
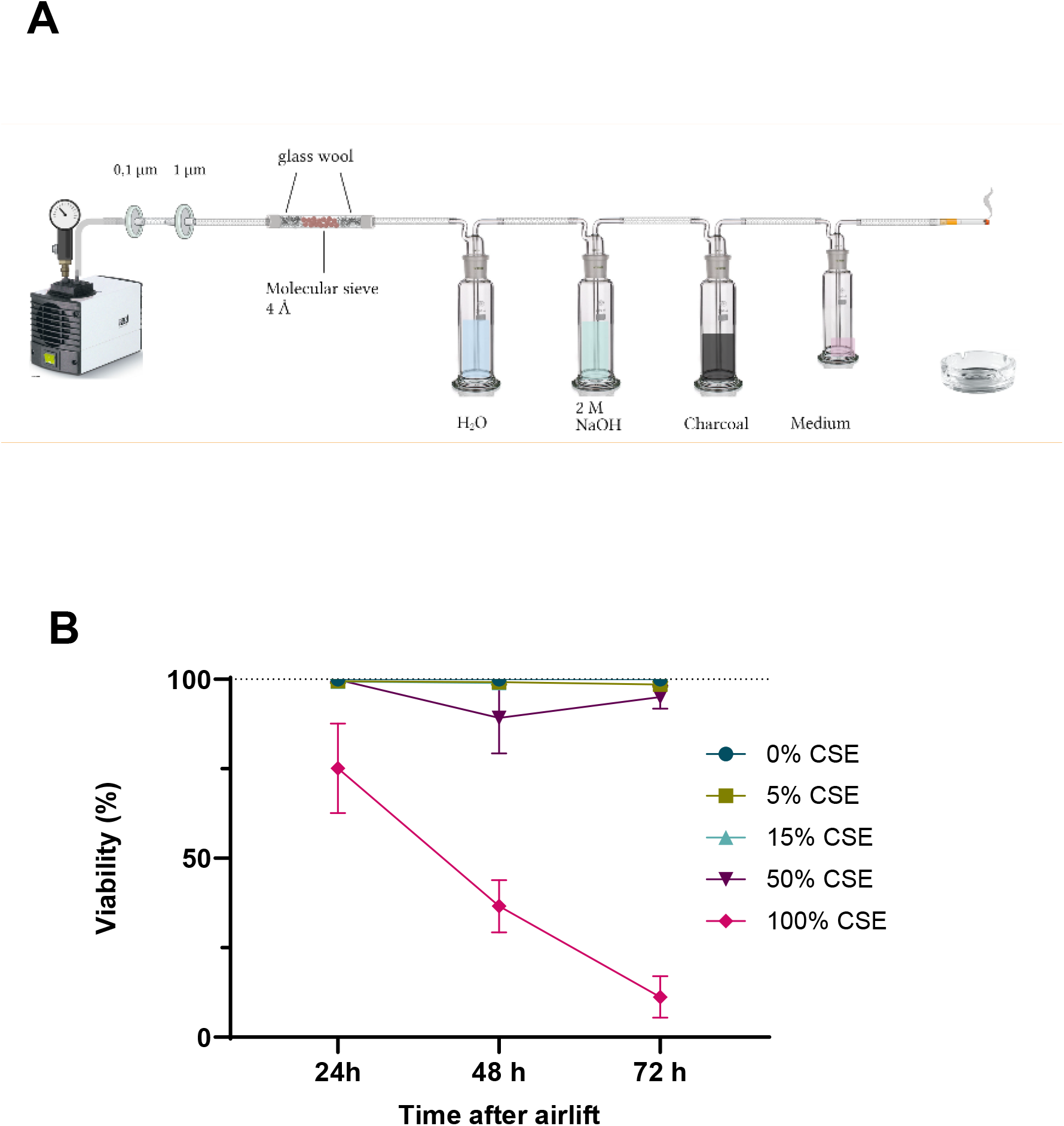
Production of Cigarette Smoke Extract (CSE) and viability of pHBECs after application of different concentrations of CSE. **A** Experimental setup for the production of 100% CSE. **B** CSE was applied chronically directly after the air lift to pHBECs in the indicated concentrations and cell lysis was quantified at the specified time points of differentation at an ALI by a lactate dehydrogenase (LDH) assay.

**Fig. S2.**
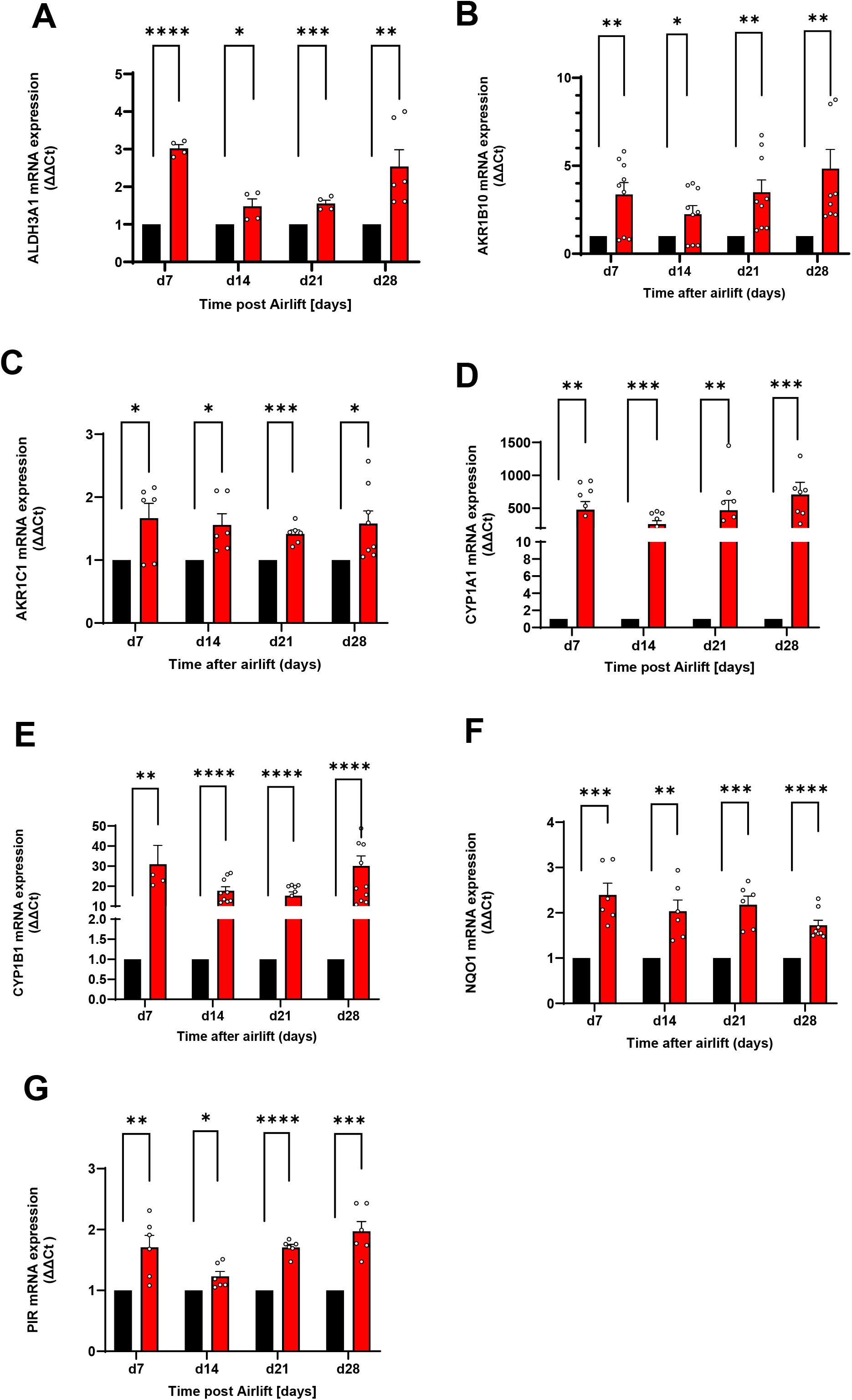
Up-regulation of mRNA expression from smoke regulated genes (SERGs) after chronic application of 5% CSE (red bars) in pHBECs from the indicated time points of differentiation at the ALI in comparison to untreated controls (black bars). Values are shown as ΔΔCt to β-actin as housekeeping gene. **A** Aldehyde dehydrogenase 3 family member A1 (ALDH3A1), **B** Aldo-keto-reductase family 1 member B10 (AKR1B10), **C** Aldo-keto-reductase family 1 member C1 (AKR1C1), **D** Cytochrome P450 1A1 (Cyp1A1), **E** Cytochrome P450 1B1 (Cyp1B1), **F** NAD(P)H quinone dehydrogenase 1 (NQO1), **G** Pirin (PIR).

**Fig. S3.**
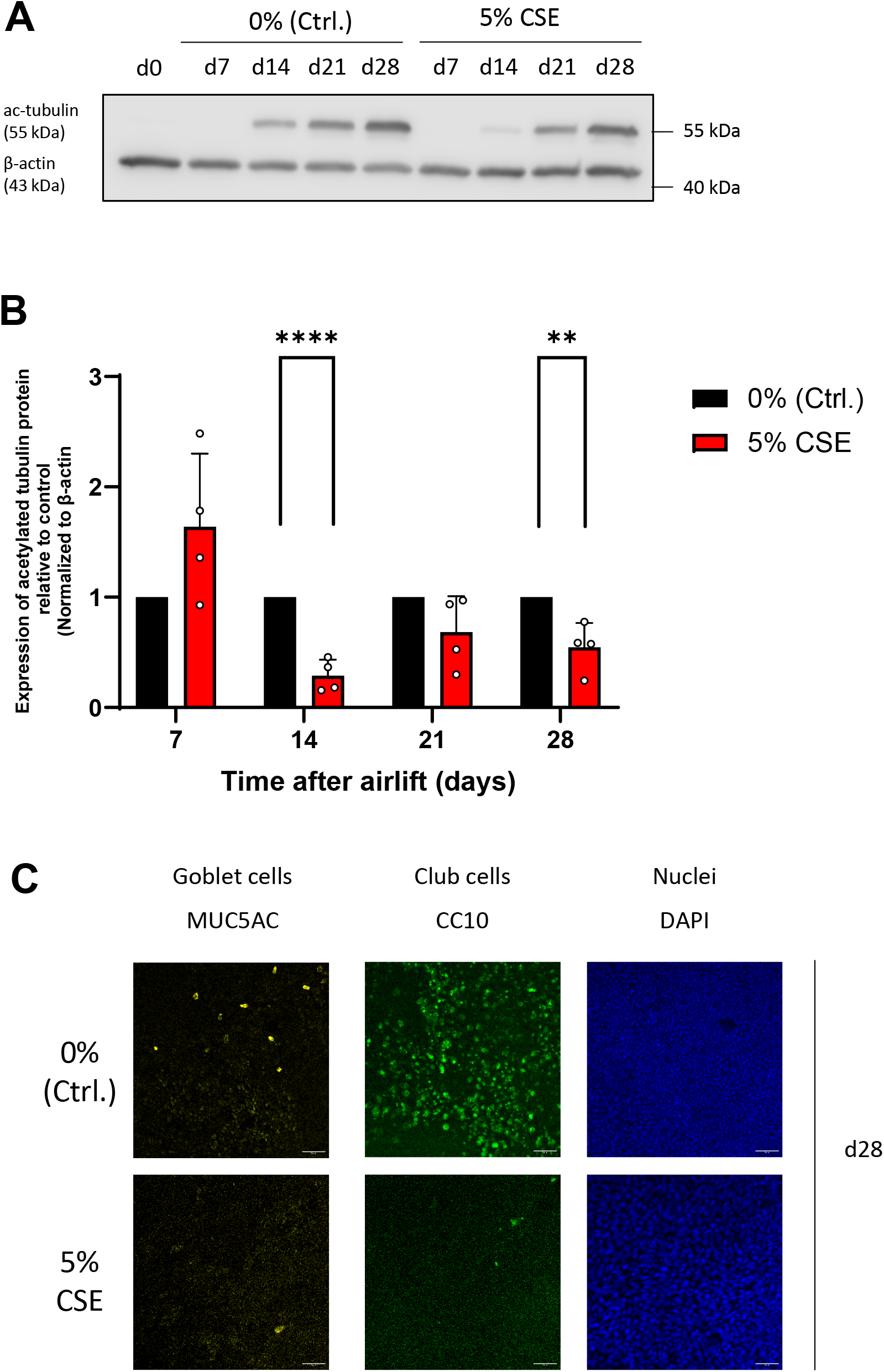
Acetylated tubulin, goblet and club cells are reduced in differentiated pHBECs after application of CSE. A Representative Western Blot of pHBECs lysates preincubatd with (5% CSE) or without CSE (0% (Ctrl.) from the indicated time points of differentiation incubated with a specific antibody directed against acetylated tubulin. Beta actin was used as loading control. B Quantification of acetylated tubulin in pHEBCs preincubated with (5% CSE, red bars) or without CSE (0% (Ctrl.), black bars). Values are shown relative to controls and normalized to β-actin expression. Data represent means + SEM of at least 4 experiments. Significance between means was analyzed using multiple unpaired Student’s t-test and is indicated as *** for p<0.001 and ** for p<0.01. C Representative confocal images of the differentiated pHBECs with (5% CSE) or without chronic CSE exposure (0%, Ctrl.). Goblet cells were identified by staining with fluorescent coupled antibodies directed against mucin 5AC (yellow, left panels) and club cells by staining with fluorescent coupled antibodies directed against Clara Cell 10-kDa protein (CC10) (green, middle panels). All cell nuclei were stained with DAPI (blue, right panels).

**Fig. S4.**
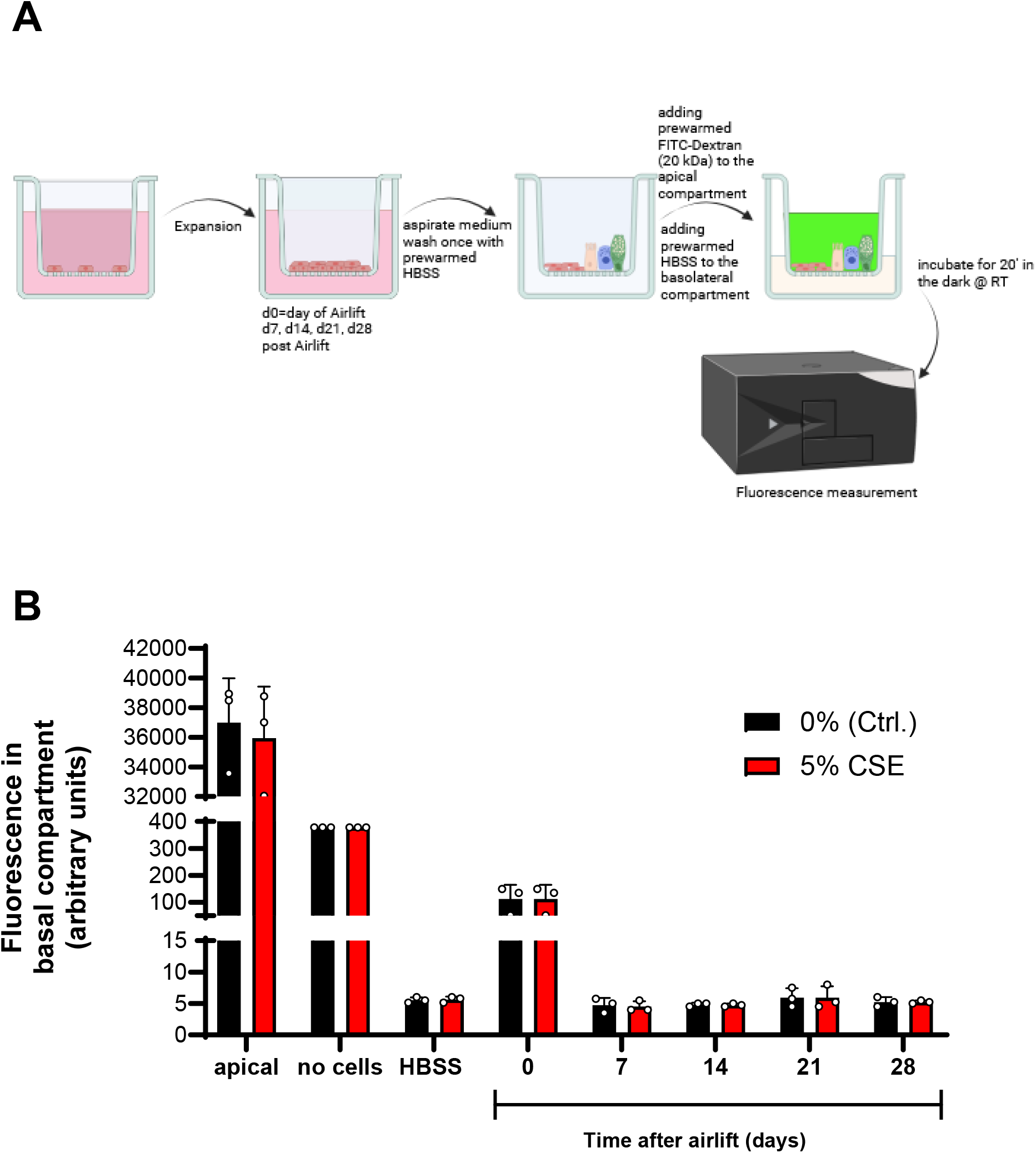
Cell permability is not changed in pHBECs differentiated at the ALI incubated with or without CSE. **A** Experimental set up of the permeability assay. After final differentiation of pHBECs a suspesion of FITC-dextrane particles were added to the apical compartment and fluorescence from the basolateral compartment was quantified in a photometer after incubation of 20 min. in the dark. **B** Quantification of cell permeability of pHBECs differentiated at the ALI incubated with (5% CSE, red bars) or without CSE (0% (Ctrl.), black bars). Fluorescence was measured in the apical (apical) or basal compartments in HBSS (HBSS), in an ALI model with no cells (no cells) or with pHBECs differentiated for 0 to 28 days.

**Fig. S5.**
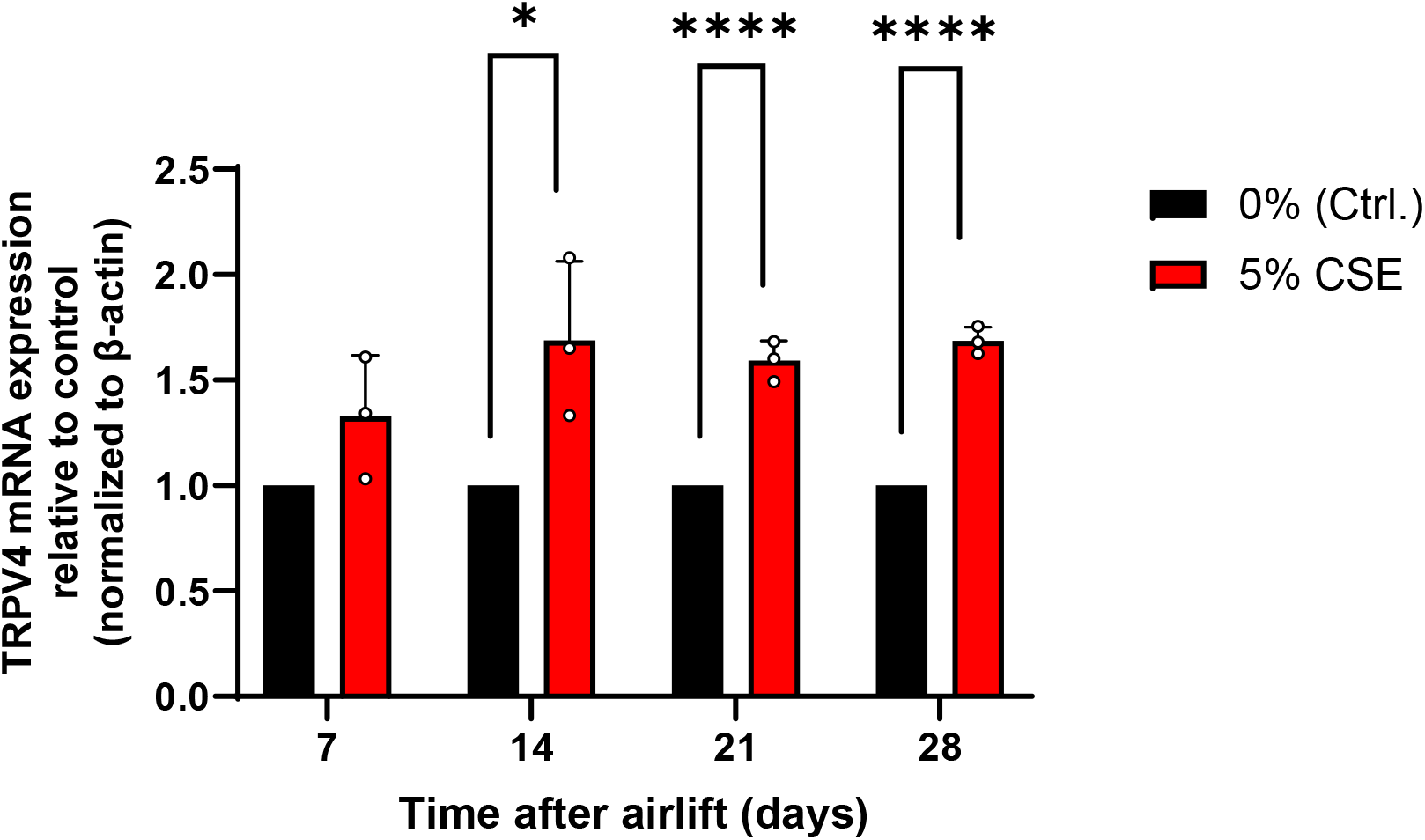
Expression of TRPV4 mRNA is increased in pHBECs differentiated at an ALI by CSE **A** Quantification of TRPV4 mRNA expression in pHBECs differentaiated for the indicated time points with (5% CSE, red bar) or without CSE (0% (Ctrl.), black bars) by quantitative reverse transcription (qRT) PCR. Values are shown relative to controls and normalized to β-actin expression. Data represent means + SEM from at least three experiments. Significance between means was analyzed using two-way ANOVA and is indicated as *** for p<0.001 and * for p<0.05.

